# HPCDb: an integrated database of pancreatic cancer

**DOI:** 10.1101/169771

**Authors:** Yonggang Tan, Yongqiang Tan, Lin Lu, Heying Zhang, Cheng Sun, Yusi Liang, Juan Zeng, Xianghong Yang, Dan Li, Huawei Zou

**Author notes:** Corresponding author: Yonggang Tan.

## Abstract

We have established a database of Human Pancreatic Cancer (HPCDb) through effectively mining, extracting, analyzing, and integrating PC-related genes, single-nucleotide polymorphisms (SNPs), and microRNAs (miRNAs), now available online at http://www.pancancer.org/. Data were extracted from established databases, ≥5 published literature (PubMed), and microarray chips (screening of differentially expressed genes using limma package in R, |log_2_ fold change (FC)| > 1). Further, protein–protein interactions (PPIs) were investigated through the Human Protein Reference Database. miRNA–target relationships were also identified using the online software TargetScan. Currently, HPCDb contains 3284 genes, 120 miRNAs, 589 SNPs, 10,139 PPIs, and 3904 miRNA–target pairs. The detailed information on PC-related genes (e.g., gene identifier (ID), symbol, synonyms, full name, chip sets, expression alteration, PubMed ID, and PPIs), miRNAs (e.g., accession number, chromosome location, related disease, PubMed ID, and miRNA–target interactions), and SNPs (e.g., SNP ID, allele, gene, PubMed ID, chromosome location, and disease) is presented through user-friendly query interfaces or convenient links to NCBI GEO, NCBI PubMed, NCBI Gene, NCBI dbSNP, and miRBase. Overall, HPCDb provides biologists with relevant information on human PC-related molecules at multiple levels, helping to generate new hypotheses or identify candidate markers.

---

As the fourth leading cause of cancer-related death and the fifth most aggressive malignancy (Nagpal et al. 2014), pancreatic cancer (PC) is estimated to cause 227,000 deaths per year globally, and its incidence and mortality rate have been gradually rising (Vincent et al. 2011). Risk factors for PC include environmental factors (e.g., insecticides, chlorinated hydrocarbon solvents, nickel, and gasoline and related compounds), medical/surgical factors (e.g., cirrhosis, chronic pancreatitis, and diabetes), genetic factors, age, sex, and lifestyle (e.g., smoking and diet) (Vincent et al. 2011). As the symptoms of PC are nonspecific, including jaundice, fatigue, loss of appetite and weight, glucose intolerance, pain in the abdomen and back, diarrhea, dizziness, chills, and muscle spasms, its early diagnosis mainly depends on biomarkers (Nolen et al. 2014). Despite substantial progress in the detection and clinical treatment of PC, only approximately 6% of patients are alive 5 years after diagnosis (Siegel et al. 2014). Moreover, PC responds poorly to chemotherapy or radiotherapy, requiring a clearer understanding of the biological mechanisms of PC and identification of its novel biomarkers (Vincent et al. 2011).

During the past decades, basic molecular experiments and high-throughput omics studies have been widely conducted to investigate the molecular mechanisms of PC, and substantial progress has been made. However, the obtained data are too scattered across the literature to be useful to biologists and pharmacologists working on PC. Moreover, some of the data derived from high-throughput omics studies have not been annotated correctly or completely. To overcome these obstacles, a few databases of PC have been established, but they exhibit both advantages and disadvantages. Pancreatic Expression Database (PED, http://www.pancreasexpression.org) is a major component of the European Union project MolDiag-Paca (Cutts et al. 2011a), which focuses mainly on pancreatic-derived omics data (e.g., genomics, transcriptomics, proteomics, and microRNAs (miRNAs)), annotations, and clinical data. Pancreatic Cancer Database (PCD, http://pancreaticcancerdatabase.org/) is a recently established database, which emphasizes literature mining for experimentally demonstrated quantitative alterations of miRNAs, mRNAs, and proteins (Thomas et al. 2014). Pancreatic Cancer Gene Database (PC-GDB, http://www.bioinformatics.org/pcgdb/) mainly provides information on genes involved in PC. Pancreatic Cancer Methylation Database (PCMDB, http://crdd.osdd.net/raghava/pcmdb/) is dedicated to the manual collection and compilation of the methylation status of genes in PC from the published literature (Nagpal et al. 2014). However, in these databases, there is little focus on PC-related single-nucleotide polymorphisms (SNPs), miRNA–target relationships, and protein–protein interactions (PPIs), which are supposed to play critical roles in cancer progression (Erichsen and Chanock 2004; Zhu et al. 2013; Sun et al. 2014).

SNPs occur every 1,000–2,000 bases (on average) in human chromosomes (Wang et al. 1998), representing the most widespread type of sequence variation in the genome (Fareed and Afzal 2013). Missense SNPs that fall within coding regions of genes may change the amino acid sequence of the corresponding proteins, while SNPs that fall within non-coding regions of genes may affect the sequence of non-coding RNA, transcription factor binding, mRNA degradation, or gene splicing, influencing individual responses to the environment, risk of disease, and prognosis (Sachidanandam et al. 2001). Thus, SNPs are valuable genetic markers for disease prevention, detection, and cure. SNPs of various genes, such as *NR5A2*, *HNF1A*, *DPYD*, *SERPINA3*, and *ABCG2*, have been identified to be associated with PC (Petersen et al. 2010; Pierce and Ahsan 2011; Zeng et al. 2011). Hence, SNP data are an important resource for PC research. Consisting of approximately 22 nucleotides, miRNAs have been reported to regulate protein-coding genes endogenously at the post-transcriptional level (Bartel 2004). Evidence has indicated that miRNAs play critical roles in PC progression through regulating their target genes, and they can serve as bio-markers for PC diagnosis, targets for clinical therapies, and criteria in prognosis prediction (Giovannetti et al. 2010; Ali et al. 2011; Srivastava et al. 2011; Liu et al. 2012; Li et al. 2013). Hence, miRNA–target data are an important resource for PC research as well. In addition, investigation of PPIs can provide novel targets for clinical therapies (Agharkar et al. 2011), suggesting the need to compile PPI information of PC-related genes.

In this study, to facilitate the systematic study of PC, HPCDb (Database for Pancreatic Cancer, http://www.pancancer.org/) was constructed through effective mining, extracting, analyzing, and integrating PC-related data of genes, SNPs, miRNAs, PPIs, and miRNA–target relationships.

## RESULTS

HPCDb provides a search engine to survey detailed information on PC-related genes, miRNAs, and SNPs, whose identifier (ID) and/or symbol could serve as query keywords. A rough flowchart of a query is presented in Figure 1.

**Figure 1.**
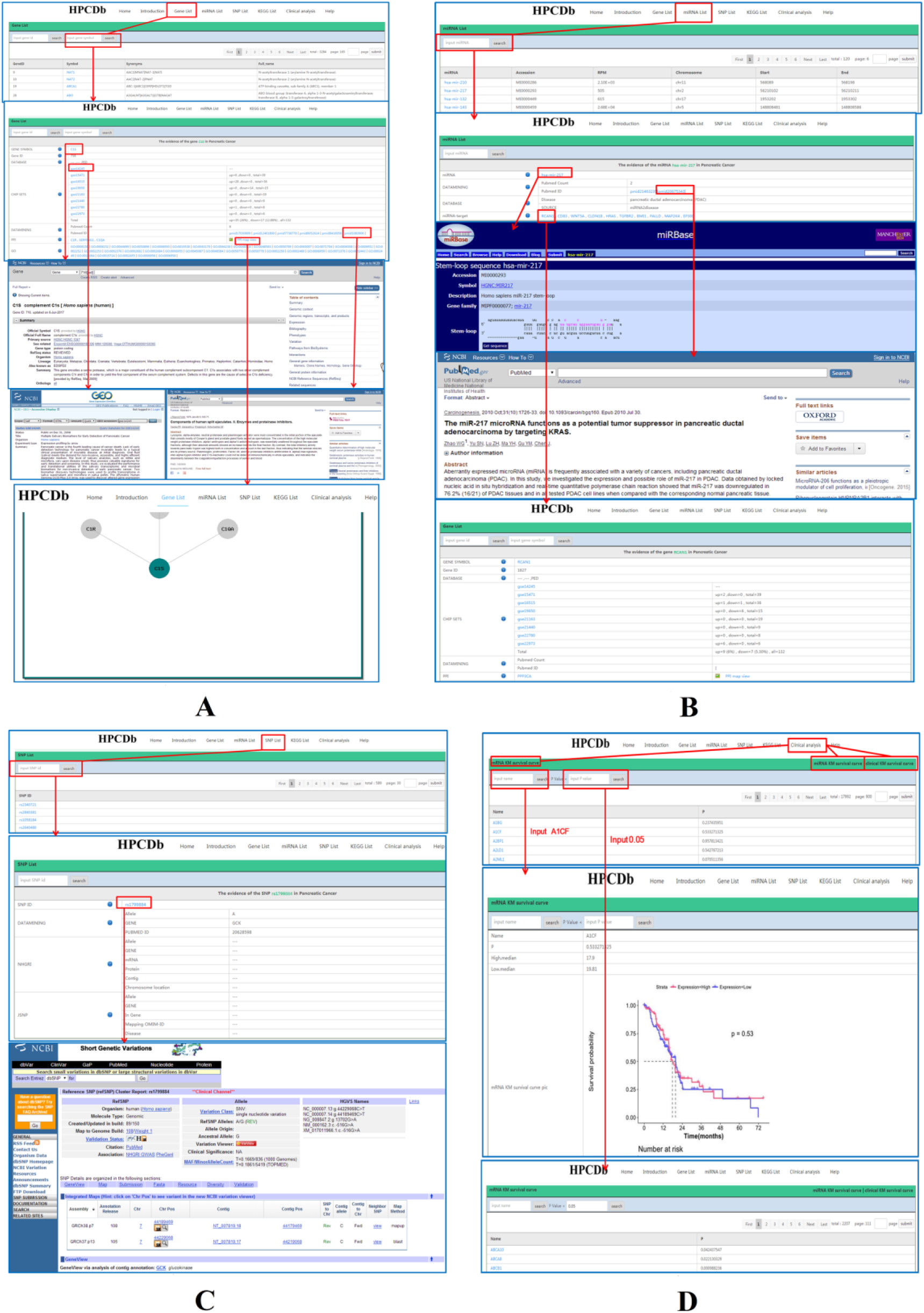
Flow charts of a query. A: A flowchart of a query on the Gene List page. B: A flowchart of a query on the miRNA List page. C: A flowchart of a query on the SNP List page. Red box and arrow: click or tap a search term and then browse; purple box and text: text in purple illustrates the term in the purple box. D: A flowchart of a query on the Clinical Analysis page.

On the “Gene List” page, PC-related genes are listed alphabetically. After submission of a gene ID or symbol, gene-centered information is displayed on a new page, including gene ID, symbol, synonyms, full name, databases featuring the present gene (PC-GDB, Ensembl, and/or PED), expression alterations based on microarray data, PubMed IDs of literature concerning the present gene, and PPIs. Upon clicking on the gene name, a new page will be displayed, where in the “chip sets” section, the word “total” demonstrates the total numbers of patients who have shown upregulation and/or downregulation of the particular gene. A further click on terms in blue will provide the link to the pages with the corresponding information in the NCBI Gene, NCBI GEO, NCBI PubMed, or HPCDb database, describing the corresponding terms in detail. A click on “PPI map view” will display a new page that illustrates the PPI network of the particular gene and the genes with which it interacts.

On the “miRNA List” page, PC-related miRNAs are also listed alphabetically. After submission of an miRNA ID or symbol, miRNA-centered information will be displayed on a new page, including miRNA ID, accession number in miRBase, reads per million (RPM, a value demonstrating the expression level of miRNA), location (chromosome, start site, and end site), results of data mining (PubMed ID and literature count), as well as databases featuring the present miRNA, disease, source, and miRNA–target relationships. A click on miRNA ID in blue will further link to a new page in miRBase that describes the corresponding miRNA in detail, and a further click on the target genes in blue will subsequently link to new pages in HPCDb that describe the corresponding target genes in detail.

On the “SNP List” page, PC-related SNPs are listed alphabetically as well. After submission of SNP ID, SNP-centered information will be displayed on a new page, including SNP ID, results of data mining (allele, gene, PubMed IDs), and databases featuring the present SNP (NHGRI GWAS Catalog and/or JSNP databases). A click on SNP ID in blue will further link to a new page in NCBI dbSNP that describes the corresponding SNP in detail.

On the “clinical analysis” page, three categories, namely, mRNA survival curve, miRNA survival curve, and clinical survival curve, are included. Taking mRNA as an example, there were 17,992 mRNA KM survival curves. On the one hand, after input of the gene name, the survival curve of the related genes can be searched. For example, after inputting ABCB1, the relationship between ABCB1 expression level and survival rate can be obtained, and p = 0.00099 is displayed, indicating a significant correlation between the expression level of ABCB1 and the survival time. On the other hand, a p-value threshold can instead be input for the search. For example, a total of 2,207 results can be obtained after selecting p < 0.05. The instructions for the miRNA survival curve and clinical survival curve are the same as the instructions for the mRNA survival curve.

## Discussion

PC is a cancer with extremely high incidence and mortality rate, for which no effective clinical therapy is currently available. To provide a platform for the comprehensive study of this disease, we established HPCDb, a database for PC-related genes, miRNAs, SNPs, and clinical analyses, through effectively mining, extracting, analyzing, integrating, and annotating the existing data scattered across various databases and the literature. Information on PC-related genes (e.g., gene ID, symbol, synonyms, full name, chip sets, expression alteration, PubMed ID, and PPIs), miRNAs (e.g., accession, RPM, chromosome location, related disease, PubMed ID, and miRNA–target interactions), SNPs (e.g., SNP ID, allele, gene, PubMed ID, chromosome location, and disease), and clinical analysis was collected and integrated into this system. Currently, the database contains 3284 genes, 120 miRNAs, 589 SNPs, 10,139 PPIs, 3904 miRNA–target pairs, and 17,992 mRNA KM survival curves, as well as 457 miNRA KM survival curves, which are presented through user-friendly query interfaces. Thus, investigators can rapidly investigate whether a gene, miRNA, or SNP is involved in PC progression, and get access to the details associated with these factors.

Compared with previous databases of PC (Cutts et al. 2011b; Nagpal et al. 2014; Thomas et al. 2014), HPCDb has several salient features. (1) Based on the microarray data downloaded from the NCBI GEO database, PC-related DEGs were rescreened, and their changes in expression in association with this disease were calculated and summarized, providing comprehensive data on the possibility of such changes. (2) Various sources were utilized, including omics data, established databases, and literature in PubMed. (3) In addition to PC-related genes and miRNAs, PC-related SNPs, PPIs between PC-related genes, and miRNA–target relationships were investigated as well, expanding the systematic understanding of PC progression. (4) Clinical analysis was performed, and prognostic analysis results of PC were visualized. (5) All of the effective information about a search term is displayed on the same page, providing easy access to the molecular information at a glance. (6) Convenient links to NCBI GEO, NCBI PubMed, NCBI Gene, NCBI dbSNP, and miRBase are provided.

Overall, as a freely available web-based resource, HPCDb provides biologists and pharmacologists with relevant information on human PC-related molecules at multiple levels, helping to generate new hypotheses or identify candidate markers.

## MATERIALS AND METHODS

### Database construction

Relevant data were extracted from established databases, published literature, and microarray chips. The collected data were then analyzed, integrated, and annotated, as shown in Figure 2.

**Figure 2.**
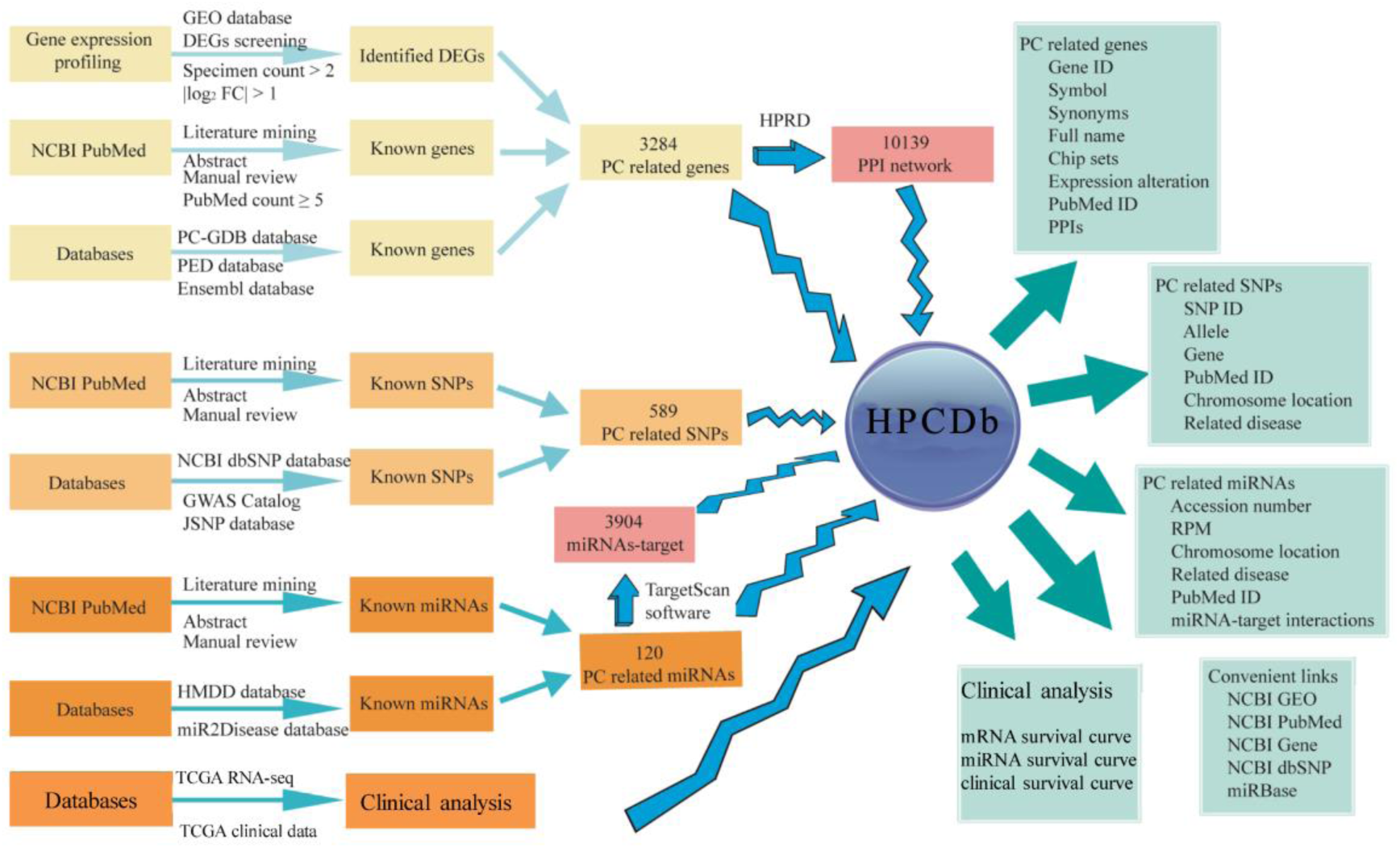
The procedure of database construction. GEO: Gene Expression Omnibus; FC: fold change; NCBI: National Center for Biotechnology Information; PC-GDB: Pancreatic Cancer Gene Database; PED: Pancreatic Expression Database; GWAS: Genome-Wide Association Studies; JSNP: Japanese Single Nucleotide Polymorphisms; HMDD: Human MicroRNA Disease Database; HPRD: Human Protein Reference Database; ID: identifier; RPM: reads per million; DEG: differentially expressed gene; PC: pancreatic cancer; PPI: protein–protein interaction; SNP: single-nucleotide polymorphism

#### PC-related genes

To collect PC-related genes, the following searches were performed: (1) known genes were extracted from PC-GDB, PED, and Ensembl (http://www.ensembl.org), which store comparative genomics, variation, and regulatory data (Flicek et al. 2012); (2) known genes with PubMed count ≥ 5 were collected; (3) gene expression profiling datasets containing samples from both a PC group (specimen count > 2) and a healthy control group were downloaded from the National Center for Biotechnology Information (NCBI) Gene Expression Omnibus (GEO, http://www.ncbi.nlm.nih.gov/geo/) database. Specifically, for each downloaded microarray dataset, raw probe-level data were preprocessed based on the corresponding platform and log-scale robust multi-array analysis (McCall et al. 2010), to conduct background correction, missing values estimation, log_2_ transformation, and value normalization. Moreover, probes without a corresponding gene symbol were omitted, and the expression values of multiple probes mapping to the same gene symbol were averaged as the gene expression value. Subsequently, the limma package in R (Smyth 2005) was utilized to identify genes that were significantly differentially expressed between PC and control groups. Only the genes with |log_2_ fold change (FC)| > 1 were defined as differentially expressed genes (DEGs), and they were also considered as PC-related genes for this database.

In total, 119 and 114 PC-related genes were extracted from PC-GDB and Ensembl, respectively, and 512 DEGs were obtained from PED. After literature mining, 1170 PC-related genes (PubMed count ≥ 5) were obtained. After analyzing gene expression data of microarray chips, 1768 significant DEGs (|log_2_ FC| > 1) were identified between the PC and control groups (Supplementary file 1). After integrating these genes, 3284 PC-related genes were finally obtained (Supplementary file 2). For these genes, Gene Ontology (GO) and Kyoto Encyclopedia of Genes and Genomes (KEGG) annotation analyses were performed to investigate their bio-functions, and the corresponding results are listed in Supplementary file 3. Further, protein–protein interactions (PPIs) of these selected genes were investigated through the Human Protein Reference Database (HPRD, http://www.hprd.org) (Keshava Prasad et al. 2009), and 10,139 PPIs were obtained (Supplementary file 4). All of the supplemented annotations of selected genes were used for database construction.

#### PC-related miRNAs

For PC-related miRNAs, the following procedures were performed: (1) known miRNAs were collected from the Human MicroRNA Disease Database (HMDD, http://www.cuilab.cn/hmdd) (Lu et al. 2008) and miR2Disease database (http://www.mir2disease.org/) (Jiang et al. 2009); and (2) known PC-related miRNAs were extracted from the literature in PubMed.

In total, 102 PC-related miRNAs were obtained from HMDD and the miR2Disease databases, whereas 38 miRNAs were obtained through literature mining. Conjunctively, 120 PC-related miRNAs were identified. Then, these miRNAs were analyzed with the online software TargetScan to predict their targets (Lewis et al. 2005; Garcia et al. 2011), from which 3904 miRNA–target pairs were identified (Supplementary file 5).

#### PC-related SNPs

For PC-related SNPs, we utilized the following search methods: (1) known SNPs were collected from the NCBI Single Nucleotide Polymorphism database (dbSNP, http://www.ncbi.nlm.nih.gov/SNP) (Sherry et al. 2001); the National Human Genome Research Institute (NHGRI), Published Genome-Wide Association Studies (GWAS) Catalog (http://www.genome.gov/), which includes 1751 manually curated publications of 11,912 SNPs (Welter et al. 2014); and the Japanese Single Nucleotide Polymorphisms database (JSNP database, http://snp.ims.u-tokyo.ac.jp/) (Hirakawa et al. 2002); and (2) known PC-related SNPs were extracted from the literature in PubMed.

Information on SNP ID (dbSNP database), risk allele, gene symbol, mRNA, and protein was collected and standardized. A total of 346, 21, and 433 PC-related SNPs were screened from NCBI dbSNP, the NHGRI GWAS Catalog, and the JSNP databases, respectively, while 150 SNPs were extracted from more than 500 published reports (Supplementary file 6). Conjunctively, 589 PC-related SNPs were identified.

#### PC-related clinical analysis

For PC-related clinical analysis, the RNA-seq and miRNA-seq level 3 data of PC in the TCGA database (https://cancergenome.nih.gov/) and PC clinical data were downloaded. Survival curve analysis for PC was performed according to the expression level of mRNAs and miRNAs as well as demographic and clinical characteristics (age, sex, and grade). Survival analysis was conducted for 17,992 mRNAs and 457 miNRAs, and 2207 mRNAs and 29 miRNAs were found to affect the survival time of PC significantly (p < 0.05). Survival analysis for age, sex, and grade found that these three clinical indices did not significantly affect the survival time of PC.

## Acknowledgments

We thank Prof. Fengping Shan and Yuan Yuan for critical advices and fruitful discussions of the article. We also thank Wei Song and his team for technical assistance.

## Author contributions

Yonggang Tan, Yoangqiang Tan and Lin Lu conceived and designed the program. Cheng Sun, Juan Zeng and Yusi Liang performed data searching, screening, downloading and standardized processing. Heying Zhang and Dan Li performed Differential gene screening, functional analysis and Text mining analysis of these genes. Yonggang Tan and Xianghong Yang performed Up-and-downstream analysis of pancreatic cancer related genes, including miRNA-target and Protein-Protein interaction. Yongqiang Tan and Huawei Zou performed survival and prognosis analysis of genes, Yoangqiang Tan and Lin Lu conducted the pancreatic cancer database and website. Yonggang Tan, Yoangqiang Tan and Lin Lu wrote the paper with help from all authors

## Conflict of interest statement

None declared.

